# Profile of proteins involved in iron homeostasis and energy metabolism in multiple organs of leptin-deficient mice

**DOI:** 10.1101/2025.03.04.641404

**Authors:** Ana Paula Oliveira Ferreira, Jessica Monteiro Volejnik Pino, Marcio Henrique Mello da Luz, Tamirez Villas Boas Petrucci, Gabriel Orefice de Souza, Jose Donato, Kil Sun Lee

## Abstract

Obesity is a global health problem. Its impacts are even more alarming when consider associated comorbidities, which include iron deficiency. Despite the importance of iron in metabolic function, the relationship between iron deficiency and metabolic dysregulation in obesity has not been fully addressed. Here, we evaluated the proteins involved in iron homeostasis and metabolism in multiple organs of leptin-deficient obese mice, a model that allows investigation of the progression of obesity. In the liver, proteins involved in glycaemia control were altered in both 3- and 5-month-old ob/ob mice (3mOB and 5mOB), but intracellular iron storage was reduced only in 3mOB. In the serum samples, glucose and cholesterol were increased in both 3mOB and 5mOB, with no alterations in iron markers. The pattern of liver proteins and serum metabolites indicate insulin-resistance state and glycemia dysregulation. In adipose tissue, 3mOB showed increased citrate synthase levels, while 5mOB showed increased MTCO2 and reduced ATP synthase levels. Ferritin was increased only in 5mOB. In the hippocampus, one of the most affected brain regions in obesity, increased MTCO2 and reduced ATP synthase levels were observed in 3mOB, while in 5mOB, increased OXCT1, a key enzyme for ketone body utilization, was observed. Reduction of ferritin levels was observed only in 5mOB. Thus, at 5 months, when the animals develop severe obesity, iron accumulates in adipose tissue but shows a reduction in the hippocampus. Our findings suggest that iron homeostasis is disrupted in a tissue-specific manner, with altered protein profiles that indicate metabolic dysregulation in multiple organs.

## Introduction

Obesity is one of the most preventable chronic diseases, yet its prevalence continues to increase worldwide across all age groups. Obesity significantly contributes to years lived with disability. It is directly associated with musculoskeletal disorders and obstructive sleep apnea [1], [2]. The burden of comorbidities further exacerbates obesity-related problems. Cardiovascular diseases are the leading consequences of obesity, but other comorbidities such as metabolic syndrome and depression are also very common [1], [3], [4]. Obesity also increases the risk of developing certain types of cancer [5], and it is one of the modifiable risk factors that contribute to the development of dementia [6], [7]. The inflammation originated from adipose tissue is known to trigger hippocampal neurodegeneration and cognitive deficit [8].

Another important yet poorly investigated comorbidity of obesity is iron deficiency. A large number of studies have reported increased prevalence of iron deficiency in population with obesity, but precise mechanisms linking both conditions are not completely understood [9], [10], [11]. Potential contributing factors include increased consumption of energy-dense and ultra-processed food that are often deficient in micronutrients [12]. Moreover, chronic low-grade inflammation associated with obesity is known to induce hepcidin production, which promotes the degradation of ferroportin in several tissues. These changes reduce dietary iron absorption and iron recycling [13], [14]. Analysis of genetic variants using mendelian randomization approach also showed causal relationship between obesity and iron deficiency anemia [15]. Thus, the occurrence of iron deficiency in obesity can be a result of complex interaction between modern dietary habits, immune responses, hormonal changes and genetic factors.

Iron is an essential micronutrient for oxygen transport, mitochondrial fuel oxidation and energy production. Thus, its deficiency can change the pattern of fuel utilization and energy expenditure, and exacerbate diabetes and metabolic syndrome in individuals with obesity [16]. Indeed, our previous study showed that diet-induced mild iron deficiency in adult rats increased fasting glycaemia [17]. Moreover, we also observed that the higher degree of obesity was associated with lower iron status in humans [18]. This evidence indicates that better management of iron homeostasis can improve the treatment of obesity.

Liver plays a central role in the regulation of systemic iron by producing and secreting hepcidin [19]. At the cellular level, iron homeostasis is controlled by key proteins, such as ferritin that reflects intracellular iron levels [20], and ferroportin, the only protein known to play a role in secretion of intracellular iron [21]. Iron deficiency occurs when overall body iron stores are low. Alternatively, inflammation can reduce the ferroportin levels and inhibit the secretion of iron from storage tissues, such as liver and macrophages, resulting in a functional iron deficiency even without body iron depletion [22]. This pathway may be important in the context of obesity with chronic inflammation, but further studies are required.

In this study, we used a monogenic leptin-deficient obese mice (ob/ob) to evaluate the levels of intracellular iron, ferritin, and ferroportin of multiple organs. Considering that chronic alterations in iron homeostasis can affect metabolic pathways, we also evaluated the protein levels of rate-limiting enzymes of several metabolic pathways, such as phosphofructokinase (PFK) in glycolysis [23], citrate synthase (CS), which is an intermediate of TCA cycle and also a quantitative indicator of mitochondria [24], cytochrome c oxidase subunit 2 (MTCO2) and component of ATP synthase (ATP5H) as converging points of complex I and II of respiratory chain [24], [25], carbamoyl-phosphate synthase (CPS1) and pyruvate carboxylase (PC) as rate-limiting enzymes of urea cycle and gluconeogenesis respectively [26], [27] (Supplementary Figure 1).

The ob/ob mouse is the most widely used model for studying obesity at the molecular level without the confounding factor of diet. The ob/ob mice develop higher degree of obesity as they age, and can reach a weight up to three times as much as wild-type mice, making them particularly interesting for the investigation of the molecular changes that occur during the progression to severe obesity [28]. To gain insight into the time-dependent alterations in the levels of iron markers and rate-limiting enzymes, multiple organs of 3- and 5-month-old mice were collected. These multi-organ profiling can help to understand the role of iron homeostasis in metabolism within the context of obesity.

## Methods

### Animal treatment

Leptin-deficient male mice (ob/ob) on a C57BL/6 genetic background were used at 3 and 5 months of age (n = 4 for each age). As control groups, 3-month-old wild-type (wt/wt) mice (n = 4) and 5-month-old heterozygous (ob/wt) lean mice (n = 3) were used. Previous studies have demonstrated that heterozygous animals (ob/wt) exhibited a pattern of weight gain, food intake, locomotor activity, body temperature, serum insulin and glucose levels, and oxygen consumption similar to homozygous controls (wt/wt). Moreover, long term treatment of ob/wt mice with exogenous leptin did not cause significant changes in the parameters cited above, indicating that as far as one allele is functional, the heterozygous animals behave similarly to wt/wt [29], [30]. In accordance with these previous data, 5-month-old ob/wt mice showed a similar body weight to 3-month-old wt/wt mice (Supplementary Figure 2). Thus, 5-month-old ob/wt mice were a suitable lean control for this study and complied with the principles of the 3Rs in the use of animals in research. All animals had ad libitum access to food and water, and maintained under 12-hour light/dark cycle at room temperature (23 ± 2 °C). All procedures were approved by the Ethics Committee on Animal Use of the Institute of Biological Sciences at the University of São Paulo (CEUA-ICB/USP) (Protocol n° 73/2017). All experiments were conducted in accordance with the ethical guidelines adopted by the Brazilian College of Animal Experimentation (COBEA) and complied with ARRIVE (Animal Research: Reporting of In Vivo Experiments).

### Samples collection

To collect the tissue samples, euthanasia was performed by cervical dislocation and decapitation. The use of anesthesia was avoided as it can affect the glucose metabolism [31]. After the decapitation, blood was collected and centrifuged at 3500 g for 10 minutes at room temperature for serum collection. Cerebral tissue, epididymal adipose tissue, gastrocnemius muscle and liver were also collected. All samples were stored at −80 °C until use. For the brain samples, hippocampal region was punched out from the tissue slice selected based on the Mouse Brain Atlas [32].

### Preparation of tissue homogenates

Tissues were homogenized in PBS at 20 % (w/v), using TissueLyser (QIAGEN) set at 50 Hz for 5 minutes. After centrifugation for 10 minutes at 3000 g and 4 °C, supernatant was recovered and divided into two portions. One portion was immediately stored at - 20 °C for the measurement of iron concentration. To the other portion, triton x-100 and cocktail of protease inhibitors (ThermoFisher Scientific 78428) were added to a final concentration of 1 %. After incubation for 30 minutes on ice, samples were centrifuged for 5 minutes at 3000 g and 4 °C, and then, the supernatant was collected and stored at −80 °C for Western Blotting.

### Measurement of iron concentration

Iron standard solution from the Total Iron-Binding Capacity (TIBC) and Serum Iron Assay Kit (Abcam: ab239715) was diluted at 0.02, 0.04, 0.06, 0.08 and 0.10 mM. Samples (30 μL) were mixed with 30 μL of precipitation solution (1 N HCl and 10 % trichloroacetic acid) and heated for 1 hour at 95 °C. Then, samples were centrifuged at 16000 g for 10 minutes at room temperature. Supernatant (50 μL) and standard solutions were mixed with 50 μL of chromogen solution (1.5 % glycolic acid and 0.5 mM ferrozine) on a 96-well plate. After incubation for 30 minutes at 30 °C, the absorbance was measured at wavelength of 560 nm.

### Glucose quantification

The Glicose HK Liquiform kit (Labtest Ref.: 137) was used to quantify glucose on serum. Reagent 1 (pH 7.8, ATP 1 mmol/L, NAD 1 mmol/L, MgCl_2_ 10 mmol/L and NaN_3_ 7.7 mmol/L) and Reagent 2 (Hexokinase 1000 U/L, Glucose-6-phosphate dehydrogenase 6000 U/L and NaN_3_ 15 mmol/L) were mixed at ratio 4:1. On a 96-well plate, 250 µL of the mixture and 2.5 µL of samples or standard solution provided in the kit were added, and incubated for 10 minutes at 37 °C. The absorbance was measured at wavelength of 340 nm.

### Urea quantification

Serum uremia was quantified using a commercial kit (Urea CE – Labtest Ref.: 27-500). Prior the assay, 3 mL of the buffer provided in kit (100 mmol/L phosphate, pH 6.9; 312 mmol/L sodium salicylate and 16.8 mmol/L sodium nitroprusside, stabilizers and preservative) were diluted in 12 mL deionized water. Then, 250 µL of the urease solution (phosphate buffer 10 mmol/L and urease ≤ 500 kU/L and stabilizers) was added to 5 mL of the diluted buffer. 5 mL of the oxidizing solution (sodium hydroxide 2.8 mol/L and sodium hypochlorite ≤ 140 mmol/L) was diluted in 45 mL of deionized water. On a 96 well-plate, 125 µL of buffered urease was mixed with 2.5 µL of samples or standard solution (70 mg/dL of urea). After incubation for 5 minutes at 37 °C, 125 µL of the diluted oxidant was added, and incubated for additional 5 minutes. The absorbance was measured at wavelength of 600 nm.

### Triglycerides quantification

In a 96-well plate, 250 µL of the reagent (buffer 50 mmol/L; pH 7.0; 4-chlorophenol 1 – 5 mmol/L; 4-aminoantipyrine 100 – 500 mmol/L; adenosine triphosphate 0.5 - 3.0 mmol/L; lipoprotein lipase 1400 – 3000 U/L; glycerol kinase 1000 – 3000 U/L; glycerol-3-phosphate oxidase 1500 – 3500 U/L; peroxidase 3000 – 7000 U/L; magnesium ions 0.1 - 1.0 mmol /L; sodium azide 14.6 mmol/L; surfactants and stabilizers) provided in Triglycerides Liquiform kit (Labtest Ref.: 87) were added onto 2.5 μL of samples or glycerol standard solution (0.25 g/L). After incubation for 10 minutes at 37 °C, absorbance was measured at wavelength of 505 nm.

### Cholesterol quantification

Cholesterol levels were quantified using commercial kits for the Cholesterol Liquiform colorimetric assay (Labtest Ref.: 76). For HDL measurement, 2.5 μL of precipitant from the HDL Cholesterol kit (Labtest Ref.: 13) was added to 2.5 μL of serum. After vortexing, the samples were centrifuged for 15 minutes at 3500 rpm and the supernatant was collected. In a 96 well-plate, 2.5 μL of serum samples, HDL samples, and their respective standard solutions were pipetted followed by addition of 250 μL of the kit reagent (buffer ≤ 100 mmol/L, pH 7.0, phenol ≤ 24 mmol/L, sodium cholate 0.005 - 0.05 %, sodium azide 14.6 mmol/L, 4aminoanthypyrine 300 – 500 µmol/L, cholesterol esterase 250 – 1000 U/L, cholesterol oxidase 250 – 1000 U/L and peroxidase 250 – 1000 U/L, cofactor, stabilizers and surfactants). After incubation for 10 minutes at 37 °C, the absorbance was measured at wavelength of 500 nm.

### Western Blot

Samples with 20 μg of total proteins were subjected to SDS-PAGE. The fractionated proteins were transferred to the PVDF membrane, previously activated in methanol and equilibrated in transfer buffer (0.192 M Glycine, 0.025 M Tris, 20 % Methanol). The membrane was treated with 5 % BSA (bovine serum albumin) or non-fat dry milk dissolved in TBS-T (0.5 M Tris, 1.5 M NaCl and 0.1 % Tween-20) for 60 minutes and washed twice with TBS-T for 5 minutes. Then the membranes were incubated overnight at 4 °C with the primary antibodies (Abcam #ab173006 for ATP5H. GeneTex #gtx54821 for Ferroportin, and #gtx32770 for OXCT1. Cell Signaling #14309S for citrate synthase, #84510S for CPS1, #4393S for FTH, #12939S for GLUT1, #2213S for GLUT4, #4212S for NMDAR, #12746S for PFKP, #66470S for pyruvate carboxylase, and #2125S for α-tubulin. ProteinTech #55070-I-AP for MTCO2. Sigma #AV45774 for PFKL). Then, the membranes were washed for 5 minutes with TBS-T, 3 times and incubated with appropriate secondary antibodies conjugated with HRP for 1 hour at room temperature. After 5 washes of 5 minutes, membranes were incubated with Immobilon Forte Western HRP substrate (Merck WBLUF0500) and digital image was acquired using Alliance 4 mini Photodocumentation system (UVITEC Cambridge; UK). The intensity of the bands was quantified using the UVIband software (UVITEC Cambridge; UK). Ponceau S staining was used for normalization.

### Statistical analysis

The lean control and obesity groups of each age were compared using T-test with Welch correction. The groups were considered significantly different when p-value was lower than 0.05. Iron levels were compared using two-way ANOVA considering the effects of age and obesity. Graphs were plotted using GraphPad Prism 8.0.1 ver.

## Results

### Liver

Two-way ANOVA showed that hepatic iron levels were not affected by age (p = 0.08), but reduced in ob/ob group regardless of age (p = 0.006). Post-Hoc tests showed significant difference between 5-month-old control group and 3-month-old ob/ob group (p = 0.018). The comparison of ob/ob groups with respective age-matching control groups did not show significant difference (Figure 1A).

**Figure 1.**
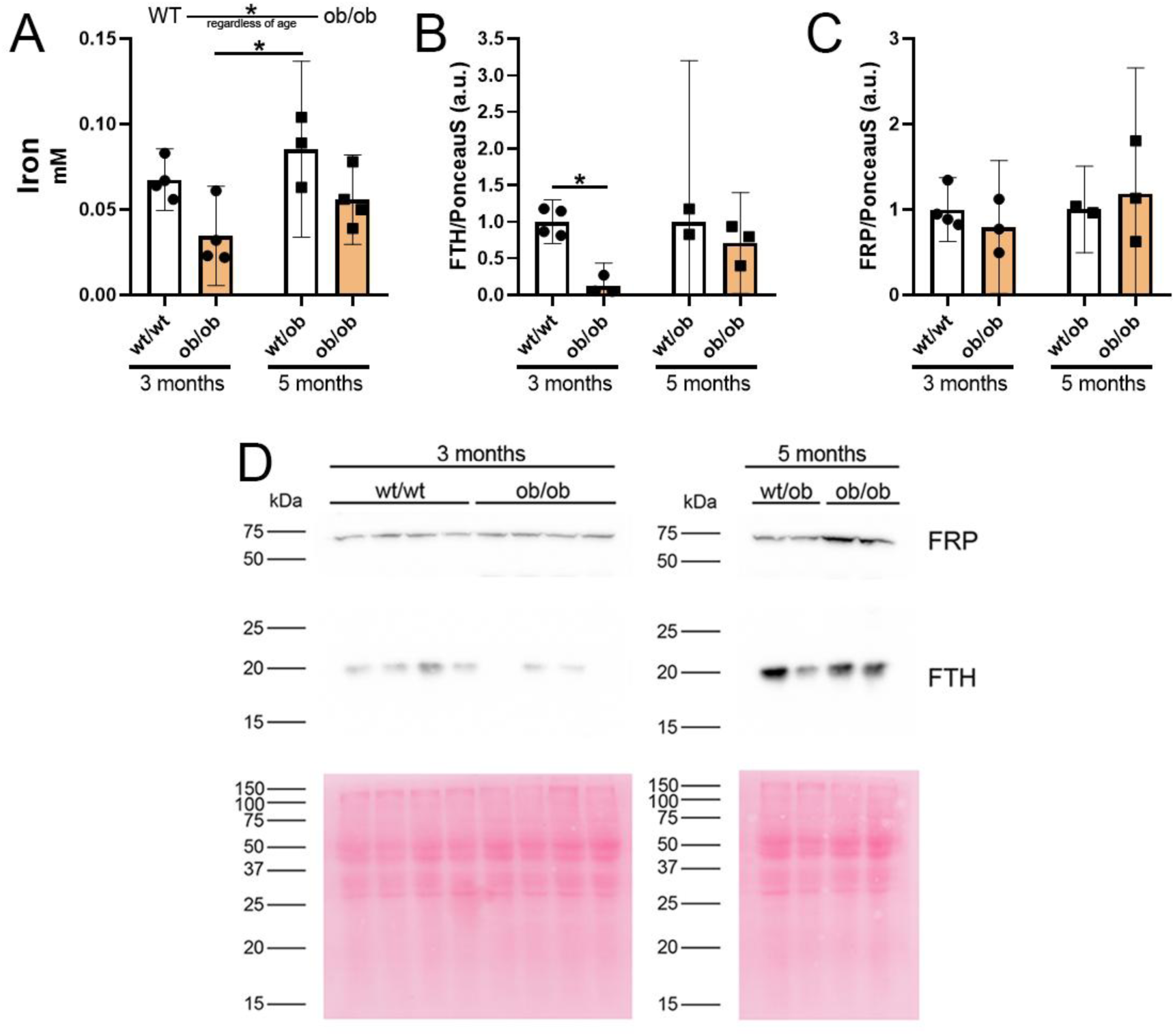
Iron levels in liver. **A**. Iron levels **B.** Quantification of ferritin (FTH) **C.** Quantification of ferroportin (FRP). The graphs were plotted by groups and age (3-month-old and 5-month-old). Bars represent the group mean, error bars represent the confidence interval of 95%, and each dot represents a biological sample. **D.** Representative images of FRP and FTH detection with respective Ponceau S. staining in 3 months-old and 5 months-old mice. * p < 0.05. Uncropped images are presented in Supplementary Figure 3.

Hepatic ferritin (FTH) levels were evaluated by western blotting and compared between groups of same age. The ferritin levels were significantly reduced in 3-month-old ob/ob mice compared to their control group, while in 5-month-old mice, no significant difference was observed (Figure 1B, middle panel of Figure 1D). Ferroportin levels were not different between the groups in both ages (Figure 1C, upper panel of Figure 1D). These results indicate that iron storage is reduced in 3-month-old ob/ob mice. This reduction is not related to iron secretion, as ferroportin levels were not altered.

Regarding the proteins involved in metabolism, we found that the level of phosphofructokinase of liver type (PFKL), one of the rate-limiting enzymes of glycolysis, was lower in both 3-month and 5-month-old animals compared to their respective controls (Figure 2A and B). The levels of citrate synthase (CS), cytochrome c oxidase subunit 2 (MTCO2), and ATP synthase were not different between the groups in both ages (Figure 2 C-H). Pyruvate carboxylase (PC), a key enzyme of the gluconeogenesis was increased in both 3-and 5-month-old obese mice (Figure 2I and J). Carbamoyl phosphate synthase (CPS1) of urea cycle was reduced in 5-month-old ob/ob mice (Figure 2K and L). These data indicate that metabolic pathways involved in glycaemia control (PFKL and PC) were affected in both 3- and 5-month-old obese mice, while amine metabolism is reduced only in 5-month-old obese mice.

**Figure 2.**
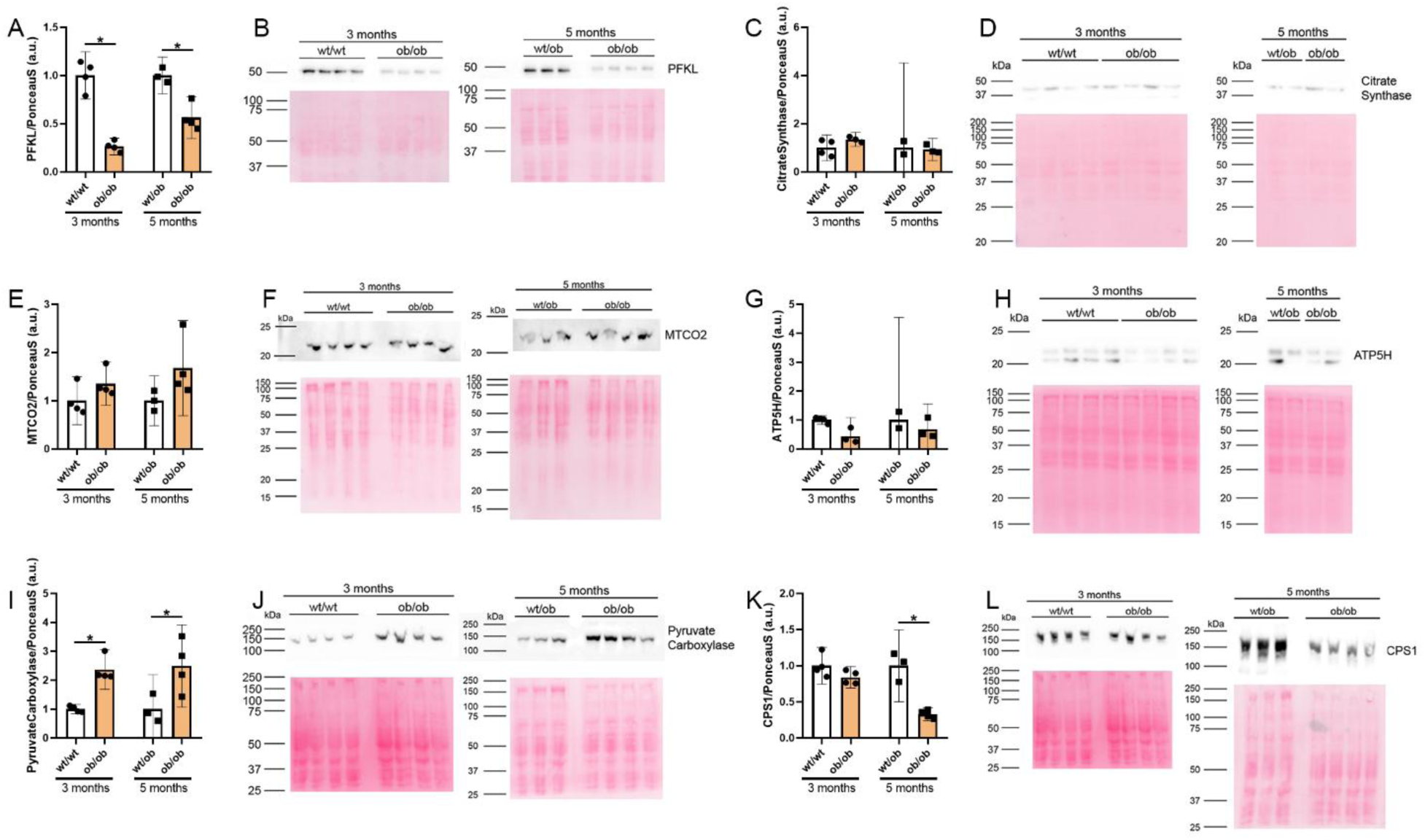
Evaluation of proteins involved in metabolism in liver. The relative quantification of liver type phosphofructokinase (PFKL, **A and B**), citrate synthase (**C and D**) mitochondrially encoded cytochrome C oxidase II (MTCO2, **E and F**), ATPase Subunit D (ATP5H, **G and H**), pyruvate carboxylase (**I and J**) and Carbamoyl phosphate synthetase 1 (CPS1, **K and L**) were carried out by western blotting. Bars represent the group mean, error bars represent the confidence interval of 95%, and each dot represents a biological sample. Representative images of immunodetection with respective ponceau S staining are presented next to each quantification. * p < 0.05. Uncropped images are presented in Supplementary Figure 4.

### Serum

In the serum samples, iron levels were not significantly altered with obesity in both ages (Figure 3A). The levels of urea were reduced only in 3-month-old mice compared to respective control (Figure 3B). Glucose levels were increased in both 3- and 5-month-old ob/ob animals compared to respective control groups (Figure 3C), while triglycerides were not different between groups (Figure 3D). Cholesterol and HDL-cholesterol were increased in ob/ob animals regardless of age (Figure 3 E and F).

**Figure 3.**
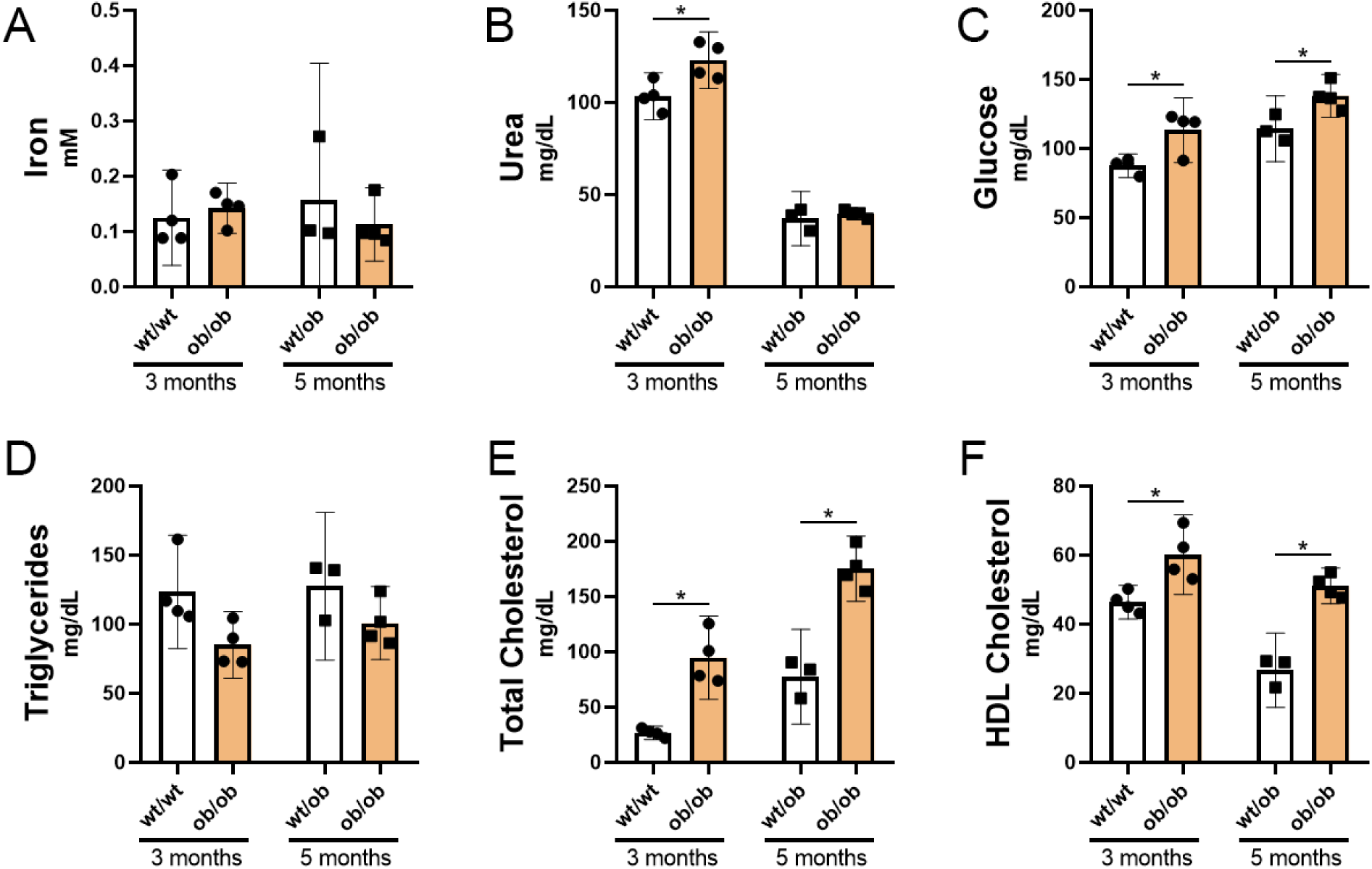
Serum metabolites. Serum levels of iron (**A**), urea (**B**), glucose (**C**), triglycerides (**D**), total cholesterol (**E**), and HDL cholesterol (**F**) were measured using colorimetric assay. Bars represent the group mean, error bars represent the confidence interval of 95 %, and each dot represents a biological sample. * p < 0.05

### Epididymal adipose tissue

One of the 3-month-old control mouse presented very little amount of adipose tissue. Thus, it was not able to confidently analyze this sample. Iron levels in epididymal adipose tissue were not statistically different between the groups in both ages (Figure 4A). However, there was a tendency for increased iron levels in 5-month-old obese mice compared to the control group. Moreover, ferritin levels were significantly increased only in 5-month-old obese mice compared to the control group (Figure 4B and C). These data indicate iron accumulation in this tissue at 5 months.

**Figure 4.**
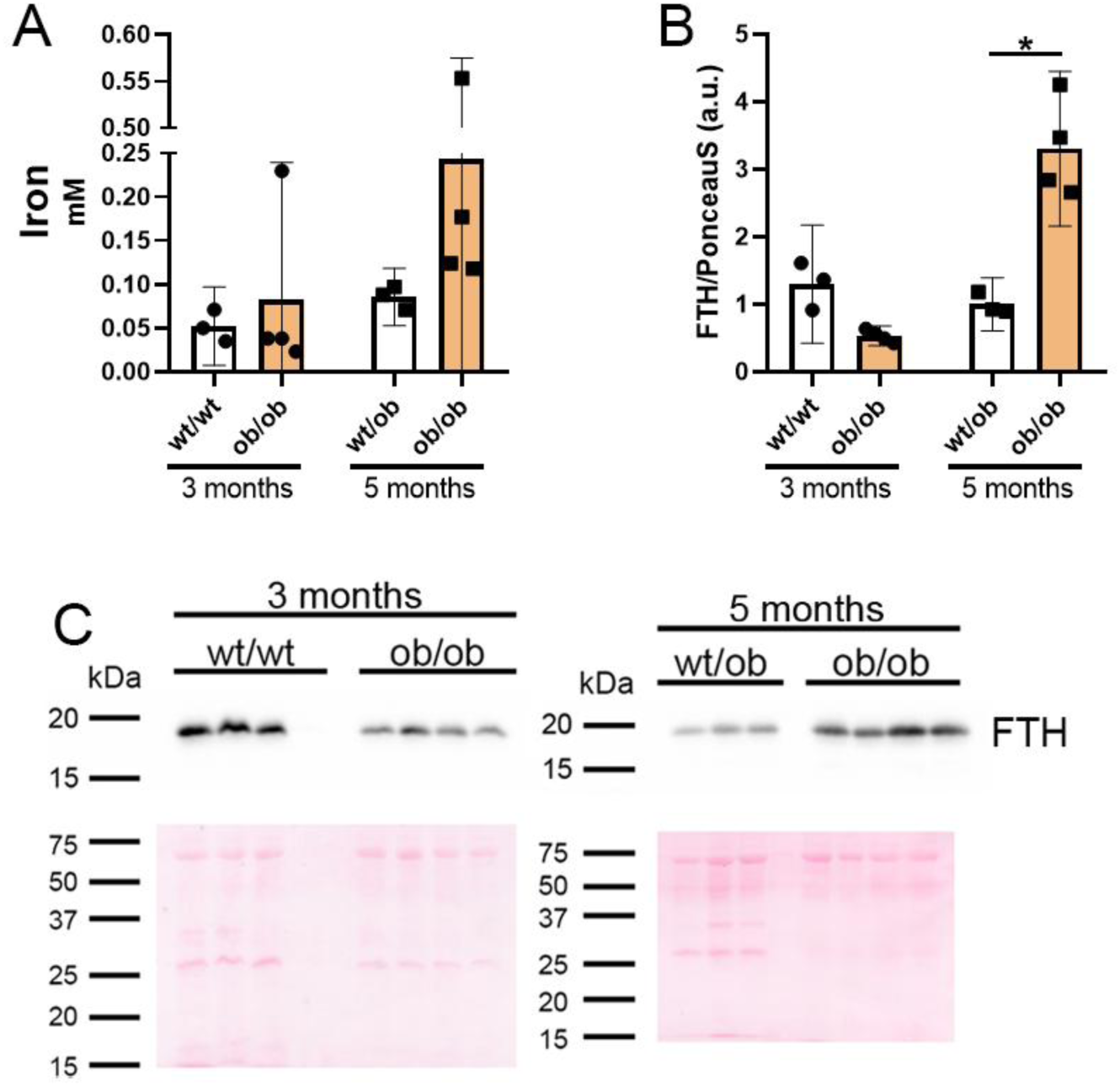
Iron levels in adipose tissue. **A**. Iron levels **B.** Quantification of ferritin (FTH). The graphs were plotted by groups and age (3-month-old and 5-month-old). Bars represent the group mean, error bars represent the confidence interval of 95 %, and each dot represents a biological sample. **C.** Representative images of FTH detection with respective Ponceau S. staining in 3 months-old and 5 months-old mice. * p < 0.05. Uncropped images are presented in Supplementary Figure 5.

The expression of GLUT4 in epididymal adipose tissue were not statistically different between groups in any age (Figure 5 A and B). Increased CS levels were observed only in 3-month-old ob/bo mice compared to their control group (Figure 5 C and D). In 5-month-old animals, MTCO2 was increased, but the expression of ATP synthase was decreased (Figure 5G and H). These non-linear alterations between citrate synthase, MTCO2 and ATP synthase indicate altered energy expenditure in this tissue at 5 months. OXCT1 which is involved in ketone body metabolism was not altered between the groups (Figure 5I and J). This result is coherent with GLUT4 levels.

**Figure 5.**
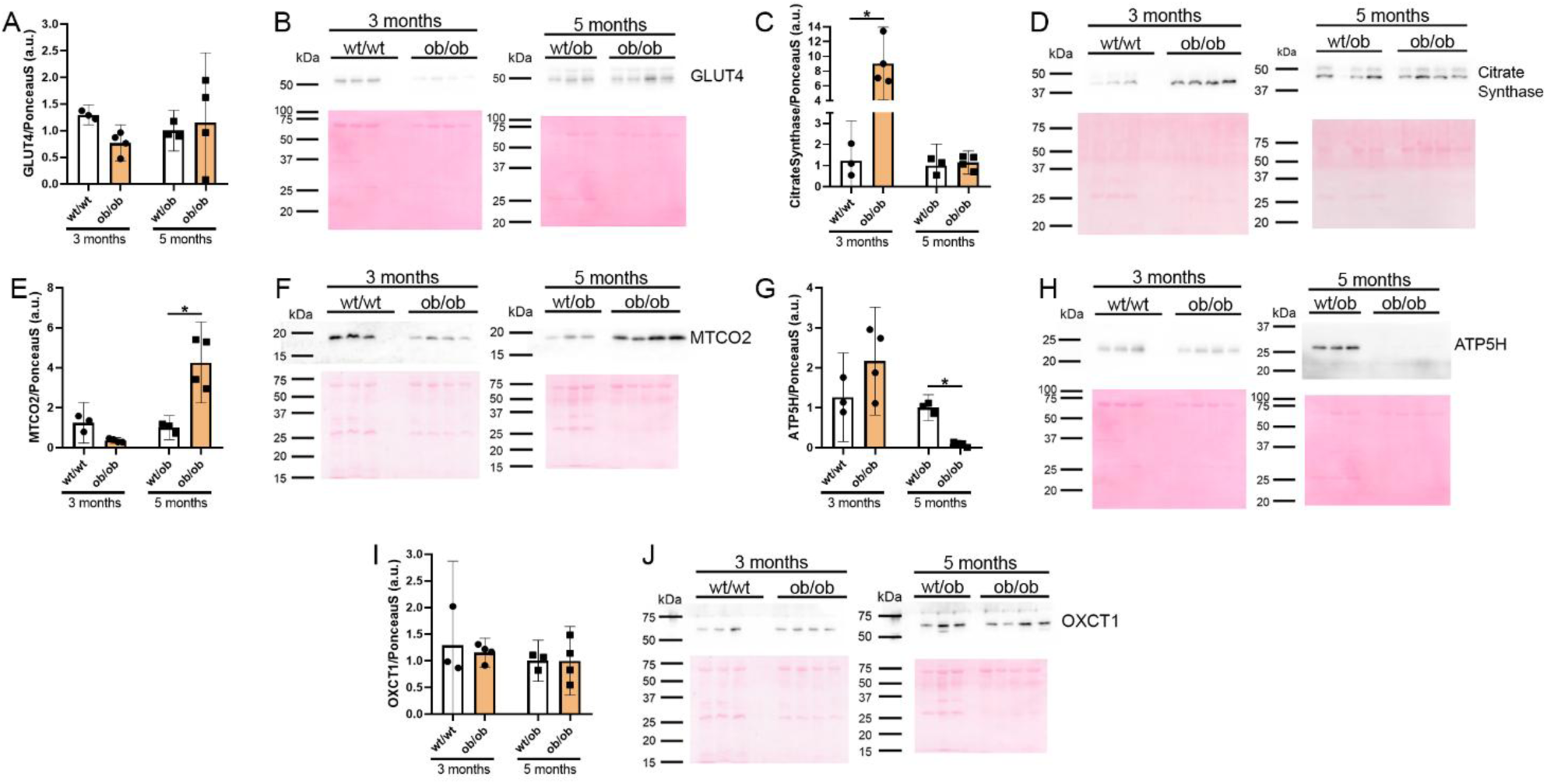
Evaluation of proteins involved in energy metabolism in epididymal adipose tissue. The relative quantification of GLUT4 (**A and B**), citrate synthase (**C and D**) mitochondrially encoded cytochrome C oxidase II (MTCO2, **E and F**), ATPase Subunit D (ATP5H, **G and H**), and 3-Oxoacid CoA-Transferase 1 (OXCT1, **I and J**) were carried out by western blotting. Bars represent the group mean, error bars represent the confidence interval of 95 %, and each dot represents a biological sample. Representative images of immunodetection with respective ponceau S staining are presented next to each quantification. * p < 0.05. Uncropped images are presented in Supplementary Figure 6.

### Gastrocnemius Muscle

Proteins involved in energy metabolism were also evaluated using gastrocnemius muscle. However, none of the proteins investigated showed significant changes between the groups (Supplementary Figure 7).

### Hippocampus

In this tissue, ferritin levels were reduced only in 5-month-old obese mice (Figure 6A and B). The levels of GLUT1 and CS were not altered between obese and control animals in both ages (Figure 6 C-F). MTCO2 was increased in 3-month-old ob/ob mice compared to control group (Figure 6G and H), while ATP synthase levels were decreased (Figure 6I and J). OXCT1 involved in ketone body catabolism was increased in the hippocampus of 5-month-old ob/ob mice (Figure 6K and L).

**Figure 6.**
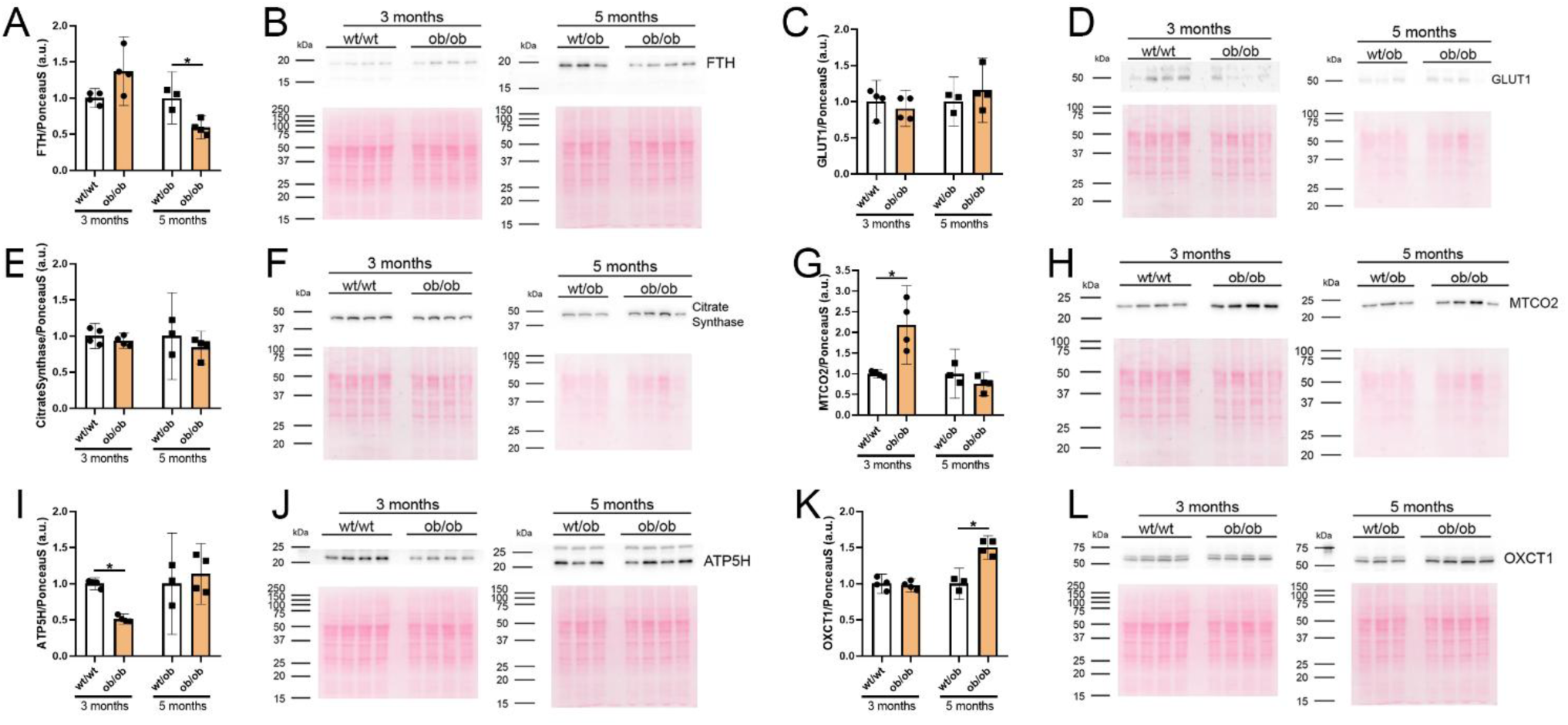
Evaluation of proteins involved in energy metabolism in hippocampus. The relative quantification of ferritin (FTH **A and B**), GLUT1 (**C and D**), citrate synthase (**E and F**), mitochondrially encoded cytochrome C oxidase II (MTCO2, **G and H**), ATPase Subunit D (ATP5H **I and J**), and 3-Oxoacid CoA-Transferase 1 (OXCT1, **K and L**) were carried out by western blotting. Bars represent the group mean, error bars represent the confidence interval of 95 %, and each dot represents a biological sample. Representative images of immunodetection with respective ponceau S staining are presented next to each quantification. * p < 0.05. Uncropped images are presented in Supplementary Figure 8.

## Discussion

To investigate the relationship between iron homeostasis and energy metabolism in the context of obesity, we evaluated the levels of the proteins involved in these processes in multiple organs of leptin-deficient obese mice at 3 and 5 months of age. The most relevant changes are summarized in Supplementary Figure 9.

In the liver, ob/ob animals of both ages showed reduced PFKL, and increased PC. These patterns suggest that the liver of obese mice behaves as if it is in a fasting state despite having free access to food [33]. These changes occurred regardless of age, while statistically significant changes in ferritin levels were observed only in 3-month-old-mice. Thus, direct relationship between liver iron levels and the impairment of glycaemia control was not observed.

Consistent with the alterations in the liver, serum glucose levels were increased in both age. These results might indicate insulin-resistance and inflammatory state that is common in obesity [34]. The increase of pyruvate carboxylase of gluconeogenesis pathway in the hepatic tissue of these animals, despite having free access to food reinforces the characteristic of an insulin-resistant state [35], [36]. Another evidence of the development of insulin-resistance in ob/ob animals is the decrease in PFKL expression on hepatocytes. This enzyme is essential for the regulation of glycolysis and its expression is controlled by insulin signaling, through the phosphorylation of fructose 2,6-bisphosphatase, which then induces the expression of PFKL [37]. In fact, insulin-resistance leads to a reduction in PFKL expression in an animal model of DM2 [38].

Once insulin-resistance is established, it is important that the animals have alternative energy sources besides glucose. The levels of CPS1, a key enzyme of the urea cycle, were assessed to investigate the degree of protein degradation, which, if exacerbated, can lead to sarcopenia, a common condition in obesity [39]. In this study, a subtle increase in serum urea was observed, but hepatic CPS1 levels remained unchanged in 3-month-old obese mice. In contrast, 5-month-old obese mice showed reduced hepatic CPS1 levels, despite no significant changes in serum urea levels. These data suggest that, by 5 months, ob/ob animals, that are severely obese, may rely less on proteins as an energy source, leading to an adaptative reduction of CPS1 levels. Additionally, OXCT1 levels were not altered in the peripheral tissues analyzed. A previous study has also found no significant changes in circulating ketone body levels between ob/ob and wt/wt animals [40], suggesting that the excessive food intake may be sufficient to meet the energy needs of peripheral tissues.

Regarding serum lipid metabolism, higher levels of total and HDL cholesterol compared to control animals are in line with data from the literature [36], [41], [42] Although HDL is increased in obese animals, the accumulation of fat in liver downregulates the expression of ABCG1 and SR-B1, preventing the proper functioning of the reverse cholesterol transport mechanism [42]. Therefore, ob/ob animals are at high risk of developing atherosclerosis and cardiovascular problems.

In adipose tissue of 3-month-old obese mice, unaltered ATP synthase and increased CS levels are apparently contradictory to data from the literature, since several studies have reported that CS activity is decreased in adipose tissue of patients with obesity [43], [44]. CS participates in the TCA cycle for energy production, but its activity is also important for de novo lipogenesis with the carbons derived from glucose [45], [46]. Thus, this molecular scenario can indicate that carbon metabolism is shifted for de novo lipogenesis. On the other hand, the iron accumulation observed in 5-month-old ob/ob mice may be related to a mechanism of adipogenesis and expansion of adipose tissue. According to the literature, iron has an important role in adipogenesis. The treatment of stem cells with iron chelator during their differentiation into adipocytes decreases the expression of adipocyte-specific genes [47]. Thus, these data may indicate that at 3-month of age, CS can contribute to the lipogenesis, while in later stages of obesity, increased iron content is necessary to generate more adipocytes. Notably, while MTCO2 was increased, ATP synthase levels were decreased in 5-month-old obese mice. Previous studies have demonstrated that increased production of reactive oxygen species by complex III and inhibition of ATP synthase promoted the differentiation of adipocytes [48], [49]. Thus, alterations in iron levels and mitochondrial proteins might play important role in hyperplasia of adipose tissue.

Hippocampus is a region of the brain known to be most vulnerable to damages caused by obesity [50] and iron deficiency [51], [52], [53]. In the hippocampus of 3-month-old obese animals, an increase in MTCO2 expression was observed with a concomitant reduction in ATP synthase levels. These data were similar to what was observed in adipose tissue of 5-month-old obese animals. While this pattern in adipose tissue may be associated with adipogenesis, in the brain, it appears to be related to mitochondrial dysfunction [54]. In the hippocampus of 5-month-old ob/ob animals, an increase in OXCT1 levels was observed, which differs from the pattern observed in peripheral tissues, but it is consistent with the insulin-resistance context and controlled lipid uptake into brain tissue.

Our findings provide insights into the temporal changes in metabolic pathways across multiple organs during the progression to severe obesity in leptin-deficient mice. Our data also indicate that iron homeostasis is dynamically altered in a tissue-specific manner. The reduction of ferritin in the hippocampus, along with its increase in adipose tissue, as well as concomitant alterations in metabolic proteins, may impact brain function and adipogenesis.

## Data availability

All data presented in this manuscript are available from the authors upon reasonable request.

## Authoŕs contributions

APOF carried out all experiments, analyzed data, and wrote the original draft of the manuscript. JMVP, MHML, TVBP, and GOS collected all tissues used in this study. JDJ helped with conceptualization of the study, and provided animals used in this study. KSL conceived the study design, supervised experiments and data analysis, and revised the manuscript. All authors read the manuscript and made intellectual contributions to the final version.

## Declaration of generative AI and AI-assisted technologies in the writing process

During the preparation of this work, the author(s) used GPT-4o mini only to check grammar and spelling.

## Ethics Statement and Patient Consent

This study did not involve human subjects.

## References

[1] A. A et al., “Health Effects of Overweight and Obesity in 195 Countries over 25 Years,” N Engl J Med, vol. 377, no. 1, pp. 13–27, Jul. 2017, doi: 10.1056/NEJMOA1614362.

[2] J. H. Lee and J. Cho, “Sleep and Obesity,” Sleep Med Clin, vol. 17, no. 1, pp. 111–116, Mar. 2022, doi: 10.1016/J.JSMC.2021.10.009.

[3] R. R. Kalyani, B. M. Everett, L. Perreault, and E. D. Michos, “Heart Disease and Diabetes,” in Diabetes in America [Internet], J. M. Lawrence, S. S. Casagrande, W. H. Herman, D. J. Wexler, and W. T. Cefalu, Eds., Bethesda (MD): National Institute of Diabetes and Digestive and Kidney Diseases (NIDDK), 2023. [Online]. Available: https://www.ncbi.nlm.nih.gov/books/NBK597416/

[4] X. Fu et al., “Shared biological mechanisms of depression and obesity: focus on adipokines and lipokines,” Aging, vol. 15, no. 12, pp. 5917–5950, 2023, doi: 10.18632/AGING.204847.

[5] S. Pati, W. Irfan, A. Jameel, S. Ahmed, and R. K. Shahid, “Obesity and Cancer: A Current Overview of Epidemiology, Pathogenesis, Outcomes, and Management,” Cancers (Basel), vol. 15, no. 2, Jan. 2023, doi: 10.3390/CANCERS15020485.

[6] G. Livingston et al., “Dementia prevention, intervention, and care: 2020 report of the Lancet Commission,” Lancet, vol. 396, no. 10248, pp. 413–446, Aug. 2020, doi: 10.1016/S0140-6736(20)30367-6.

[7] T. Dukelow et al., “Modifiable risk factors for dementia, and awareness of brain health behaviors: Results from the Five Lives Brain Health Ireland Survey (FLBHIS),” Front Psychol, vol. 13, Jan. 2023, doi: 10.3389/FPSYG.2022.1070259.

[8] T. H. Y. Lee and S. Y. Yau, “From Obesity to Hippocampal Neurodegeneration: Pathogenesis and Non-Pharmacological Interventions,” Int J Mol Sci, vol. 22, no. 1, pp. 1–33, Jan. 2020, doi: 10.3390/IJMS22010201.

[9] Z. Chen, B. Cao, L. Liu, X. Tang, and H. Xu, “Association between obesity and anemia in an nationally representative sample of United States adults: a cross-sectional study,” Front Nutr, vol. 11, 2024, doi: 10.3389/FNUT.2024.1304127.

[10] P. N. Benotti et al., “Iron deficiency is highly prevalent among candidates for metabolic surgery and may affect perioperative outcomes,” Surg Obes Relat Dis, vol. 17, no. 10, pp. 1692–1699, Oct. 2021, doi: 10.1016/J.SOARD.2021.05.034.

[11] L. Zhao, X. Zhang, Y. Shen, X. Fang, Y. Wang, and F. Wang, “Obesity and iron deficiency: a quantitative meta-analysis,” Obes Rev, vol. 16, no. 12, pp. 1081–1093, Dec. 2015, doi: 10.1111/OBR.12323.

[12] V. S. Malik, W. C. Willett, and F. B. Hu, “Global obesity: trends, risk factors and policy implications,” Nat Rev Endocrinol, vol. 9, no. 1, pp. 13–27, Jan. 2013, doi: 10.1038/NRENDO.2012.199.

[13] A. C. Cepeda-Lopez and K. Baye, “Obesity, iron deficiency and anaemia: a complex relationship,” Public Health Nutr, vol. 23, no. 10, pp. 1703–1704, Jul. 2020, doi: 10.1017/S1368980019004981.

[14] N. U. Stoffel et al., “The effect of central obesity on inflammation, hepcidin, and iron metabolism in young women,” Int J Obes (Lond), vol. 44, no. 6, pp. 1291–1300, Jun. 2020, doi: 10.1038/S41366-020-0522-X.

[15] T. Wang et al., “Causal relationship between obesity and iron deficiency anemia: a two-sample Mendelian randomization study,” Front Public Health, vol. 11, 2023, doi: 10.3389/FPUBH.2023.1188246.

[16] S. L. Kim, S. Shin, and S. J. Yang, “Iron Homeostasis and Energy Metabolism in Obesity,” Clin Nutr Res, vol. 11, no. 4, p. 316, 2022, doi: 10.7762/CNR.2022.11.4.316.

[17] J. M. V. Pino et al., “Iron-deficient diet induces distinct protein profile related to energy metabolism in the striatum and hippocampus of adult rats,” Nutr Neurosci, vol. 25, no. 2, pp. 1–12, 2022, doi: 10.1080/1028415X.2020.1740862.

[18] J. M. V. Pino et al., “Severe Obesity in Women Can Lead to Worse Memory Function and Iron Dyshomeostasis Compared to Lower Grade Obesity,” Int J Endocrinol, vol. 2023, 2023, doi: 10.1155/2023/7625720.

[19] C. Y. Wang and J. L. Babitt, “Liver iron sensing and body iron homeostasis,” Blood, vol. 133, no. 1, pp. 18–29, Jan. 2019, doi: 10.1182/BLOOD-2018-06-815894.

[20] F. M. Torti and S. V Torti, “Regulation of ferritin genes and protein,” Blood, vol. 99, no. 10, pp. 3505–3516, May 2002, doi: 10.1182/blood.v99.10.3505.

[21] E. Nemeth and T. Ganz, “Hepcidin-ferroportin interaction controls systemic iron homeostasis,” Jun. 02, 2021, MDPI. doi: 10.3390/ijms22126493.

[22] S. R. Pasricha, J. Tye-Din, M. U. Muckenthaler, and D. W. Swinkels, “Iron deficiency,” Lancet, vol. 397, no. 10270, pp. 233–248, Jan. 2021, doi: 10.1016/S0140-6736(20)32594-0.

[23] I. Mor, E. C. Cheung, and K. H. Vousden, “Control of glycolysis through regulation of PFK1: Old friends and recent additions,” Cold Spring Harb Symp Quant Biol, vol. 76, pp. 211–216, 2011, doi: 10.1101/sqb.2011.76.010868.

[24] J. M. Małecki, E. Davydova, and P. Falnes, “Protein methylation in mitochondria,” Apr. 01, 2022, American Society for Biochemistry and Molecular Biology Inc. doi: 10.1016/j.jbc.2022.101791.

[25] J. S. Yook et al., “Dietary iron deficiency modulates adipocyte iron homeostasis, adaptive thermogenesis, and obesity in C57BL/6 mice,” Journal of Nutrition, vol. 151, no. 10, pp. 2967–2975, Oct. 2021, doi: 10.1093/jn/nxab222.

[26] C. Diez-Fernandez and J. Häberle, “Targeting CPS1 in the treatment of Carbamoyl phosphate synthetase 1 (CPS1) deficiency, a urea cycle disorder,” Apr. 03, 2017, Taylor and Francis Ltd. doi: 10.1080/14728222.2017.1294685.

[27] N. Kumashiro et al., “Targeting pyruvate carboxylase reduces gluconeogenesis and adiposity and improves insulin resistance,” Diabetes, vol. 62, no. 7, pp. 2183–2194, Jul. 2013, doi: 10.2337/db12-1311.

[28] J. B. Suleiman, M. Mohamed, and A. B. A. Bakar, “A systematic review on different models of inducing obesity in animals: Advantages and limitations,” J Adv Vet Anim Res, vol. 7, no. 1, pp. 103–114, Mar. 2019, doi: 10.5455/JAVAR.2020.G399.

[29] M. Pelleymounter et al., “Effects of the obese gene product on body weight regulation in ob/ob mice.,” Science (1979), vol. 269, no. 5223, pp. 540–543, Jul. 1995, doi: 10.1126/science.7624776.

[30] C. Tankersley et al., “Modified control of breathing in genetically obese (ob/ob) mice,” J Appl Physiol, vol. 81, no. 2, pp. 716–723, Aug. 1985, doi: 10.1152/jappl.1996.81.2.716.

[31] S. Prando, C. de G. Carneiro, D. A. Otsuki, and M. T. Sapienza, “Effects of ketamine/xylazine and isoflurane on rat brain glucose metabolism measured by 18 F-fluorodeoxyglucose-positron emission tomography,” Eur J Neurosci, vol. 49, no. 1, pp. 51–61, Jan. 2019, doi: 10.1111/EJN.14252.

[32] G. D. Rosen et al., “The Mouse Brain Library.” [Online]. Available: www.mbl.org

[33] L. Rui, “Energy metabolism in the liver,” Compr Physiol, vol. 4, no. 1, pp. 177–197, 2014, doi: 10.1002/CPHY.C130024.

[34] N. M. Leguisamo et al., “GLUT4 content decreases along with insulin resistance and high levels of inflammatory markers in rats with metabolic syndrome,” Cardiovasc Diabetol, vol. 11, Aug. 2012, doi: 10.1186/1475-2840-11-100.

[35] E. Grosbellet et al., “Leptin modulates the daily rhythmicity of blood glucose,” Chronobiol Int, vol. 32, no. 5, pp. 637–649, Jun. 2015, doi: 10.3109/07420528.2015.1035440.

[36] F. Suriano et al., “Novel insights into the genetically obese (ob/ob) and diabetic (db/db) mice: two sides of the same coin,” Microbiome, vol. 9, no. 1, Dec. 2021, doi: 10.1186/S40168-021-01097-8.

[37] P. Ausina, D. Da Silva, D. Majerowicz, P. Zancan, and M. Sola-Penna, “Insulin specifically regulates expression of liver and muscle phosphofructokinase isoforms,” Biomed Pharmacother, vol. 103, pp. 228–233, Jul. 2018, doi: 10.1016/J.BIOPHA.2018.04.033.

[38] K. Dong, H. Ni, M. Wu, Z. Tang, M. Halim, and D. Shi, “ROS-mediated glucose metabolic reprogram induces insulin resistance in type 2 diabetes,” Biochem Biophys Res Commun, vol. 476, no. 4, pp. 204–211, Aug. 2016, doi: 10.1016/J.BBRC.2016.05.087.

[39] L. P. C. Mayoral et al., “Obesity subtypes, related biomarkers & heterogeneity,” Indian J Med Res, vol. 151, no. 1, pp. 11–21, Jan. 2020, doi: 10.4103/IJMR.IJMR_1768_17.

[40] P. Giesbertz et al., “Metabolite profiling in plasma and tissues of ob/ob and db/db mice identifies novel markers of obesity and type 2 diabetes,” Diabetologia, vol. 58, no. 9, pp. 2133–2143, Sep. 2015, doi: 10.1007/S00125-015-3656-Y.

[41] L. Arisqueta, H. Navarro-Imaz, Y. Rueda, and O. Fresnedo, “Cholesterol mobilization from hepatic lipid droplets during endotoxemia is altered in obese ob/ob mice,” J Biochem, vol. 158, no. 4, pp. 321–329, Mar. 2015, doi: 10.1093/JB/MVV047.

[42] M. N. Duong, K. Uno, V. Nankivell, C. Bursill, and S. J. Nicholls, “Induction of obesity impairs reverse cholesterol transport in ob/ob mice,” PLoS One, vol. 13, no. 9, Sep. 2018, doi: 10.1371/JOURNAL.PONE.0202102.

[43] M. Christe et al., “Obesity affects mitochondrial citrate synthase in human omental adipose tissue,” ISRN Obes, vol. 2013, pp. 1–8, Jul. 2013, doi: 10.1155/2013/826027.

[44] Y. X. Xiao, I. R. Lanza, J. M. Swain, M. G. Sarr, K. S. Nair, and M. D. Jensen, “Adipocyte mitochondrial function is reduced in human obesity independent of fat cell size,” J Clin Endocrinol Metab, vol. 99, no. 2, Feb. 2014, doi: 10.1210/JC.2013-3042.

[45] W. Liu et al., “IL-1R-IRAKM-Slc25a1 signaling axis reprograms lipogenesis in adipocytes to promote diet-induced obesity in mice,” Nat Commun, vol. 13, no. 1, Dec. 2022, doi: 10.1038/S41467-022-30470-W.

[46] W. Y. Hsiao and D. A. Guertin, “De Novo Lipogenesis as a Source of Second Messengers in Adipocytes,” Curr Diab Rep, vol. 19, no. 11, Nov. 2019, doi: 10.1007/S11892-019-1264-9.

[47] D. F. Edwards et al., “Differential Iron Requirements for Osteoblast and Adipocyte Differentiation,” JBMR Plus, vol. 5, no. 9, Sep. 2021, doi: 10.1002/JBM4.10529.

[48] K. V. Tormos et al., “Mitochondrial complex III ROS regulate adipocyte differentiation,” Cell Metab, vol. 14, no. 4, pp. 537–544, Oct. 2011, doi: 10.1016/j.cmet.2011.08.007.

[49] A. Carrière, Y. Fernandez, M. Rigoulet, L. Pénicaud, and L. Casteilla, “Inhibition of preadipocyte proliferation by mitochondrial reactive oxygen species,” FEBS Lett, vol. 550, no. 1–3, pp. 163–167, Aug. 2003, doi: 10.1016/S0014-5793(03)00862-7.

[50] Y. Milaneschi, W. K. Simmons, E. F. C. van Rossum, and B. W. Penninx, “Depression and obesity: evidence of shared biological mechanisms,” Mol Psychiatry, vol. 24, no. 1, pp. 18–33, Jan. 2019, doi: 10.1038/S41380-018-0017-5.

[51] B. B. Yavuz et al., “Iron deficiency can cause cognitive impairment in geriatric patients,” J Nutr Health Aging, vol. 16, no. 3, pp. 220–224, 2012, doi: 10.1007/S12603-011-0351-7.

[52] I. Jáuregui-Lobera, “Iron deficiency and cognitive functions,” Neuropsychiatr Dis Treat, vol. 10, pp. 2087–2095, Nov. 2014, doi: 10.2147/NDT.S72491.

[53] A. Grandone, P. Marzuillo, L. Perrone, and E. M. del Giudice, “Iron Metabolism Dysregulation and Cognitive Dysfunction in Pediatric Obesity: Is There a Connection?,” Nutrients, vol. 7, no. 11, pp. 9163–9170, Nov. 2015, doi: 10.3390/NU7115458.

[54] P. Manna and S. K. Jain, “Obesity, Oxidative Stress, Adipose Tissue Dysfunction, and the Associated Health Risks: Causes and Therapeutic Strategies,” Metab Syndr Relat Disord, vol. 13, no. 10, pp. 423–444, Dec. 2015, doi: 10.1089/MET.2015.0095.

