## Supplementary figure for "Profile of proteins involved in iron homeostasis and energy metabolism in multiple organs of leptin-deficient mice"

**Supplementary Material**

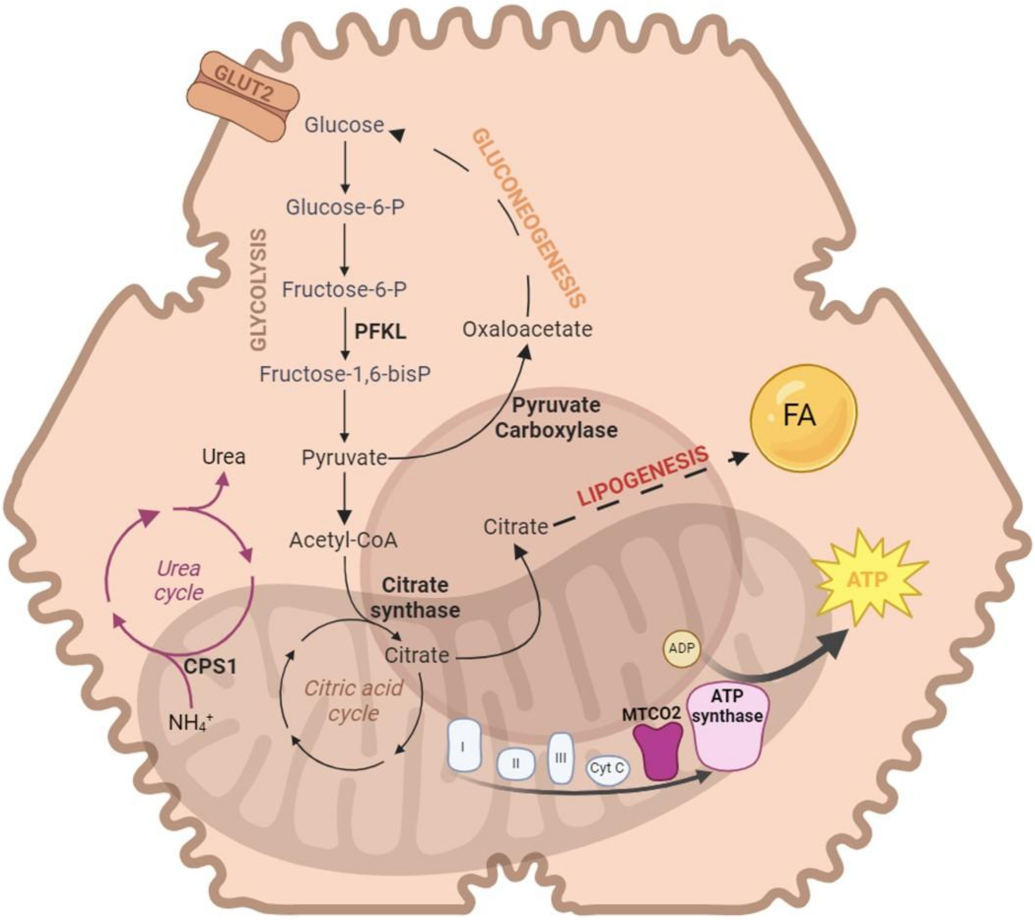


**Supplementary Figure 1. Schematic representation of metabolic pathways.** Proteins that were analyzed in this study are highlighted in bold. CPS1: carbamoyl phosphate synthetase 1, FA: fatty acids, MTCO2: mitochondrially encoded cytochrome C oxidase II, PFKL: phosphofructokinase liver type.

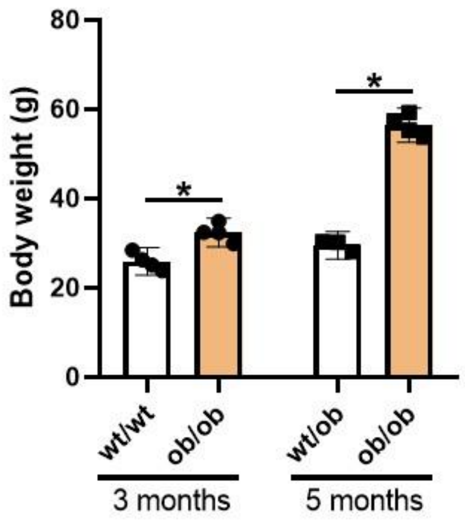


**Supplementary Figure 2. Body weight of the animals at the moment of euthanasia.** Bars represent the group mean, error bars represent the confidence interval of 95%, and each dot represents a biological sample.

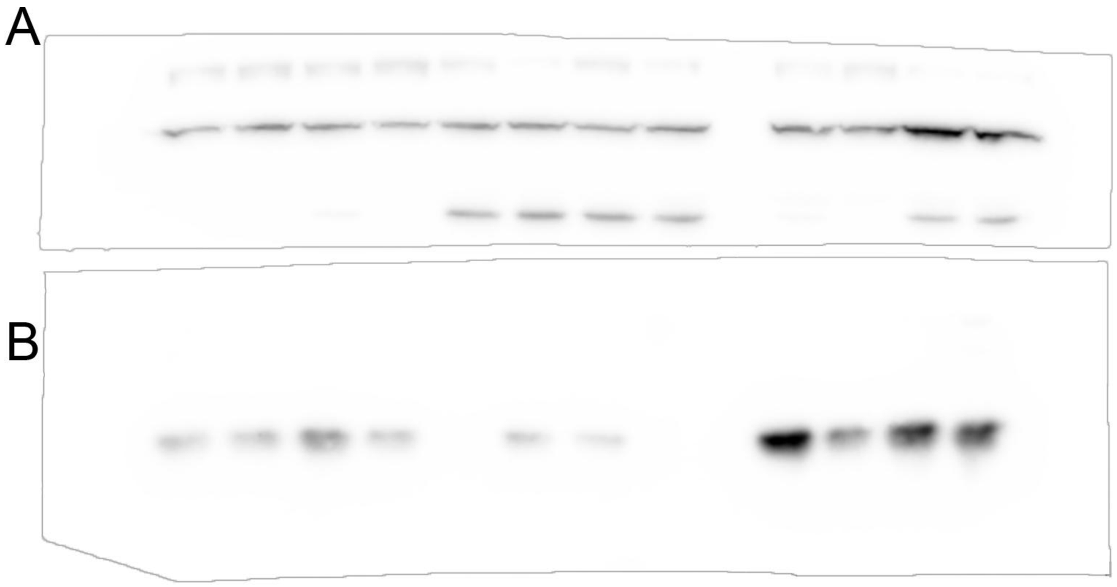


**Supplementary Figure 3. Liver ferroportin and ferritin.** Uncropped images of Figure 1. The membrane was cut at 37kDa to enable the detection of two proteins with distinct molecular weights using the same membrane. **A.** Ferroportin. **B.** Ferritin.

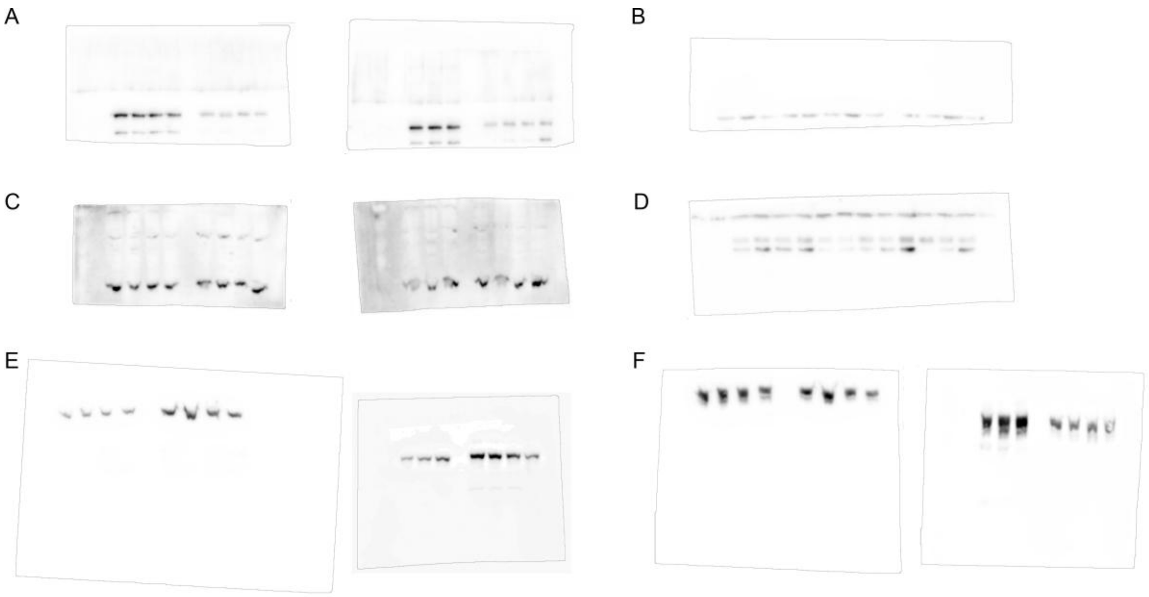


**Supplementary Figure 4. Metabolic proteins in the liver.** Uncropped images of Figure 2. **A.** PFKL. **B.** Citrate synthase. **C.** MTCO2. **D.** ATP5H. **E.** Pyruvate Carboxylase. **F.** CPS1. In some experiments, the membrane was cut at 37kDa to enable the detection of two proteins with distinct molecular weights using the same membrane.

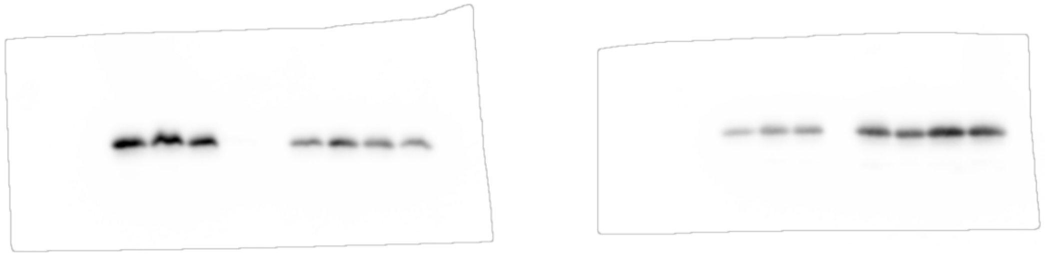

**Supplementary Figure 5. Ferritin of adipose tissue.** Uncropped images of Figure 4. Membranes were cut at 37kDa to enable the detection of two proteins with distinct molecular weights using the same membrane.

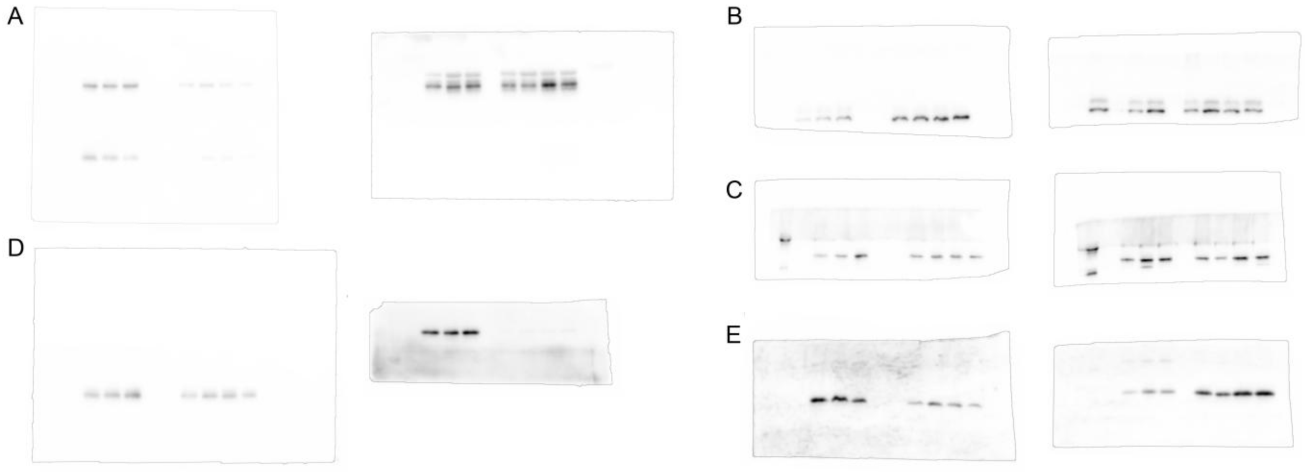


**Supplementary Figure 6. Metabolic proteins in the adipose tissue.** Uncropped images of Figure 5. **A.** GLUT4. **B.** Citrate synthase. **C.** OXCT1. **D.** ATP5H. **E.** MTCO2.

In some experiments, the membrane was cut at 37kDa to enable the detection of two proteins with distinct molecular weights using the same membrane.

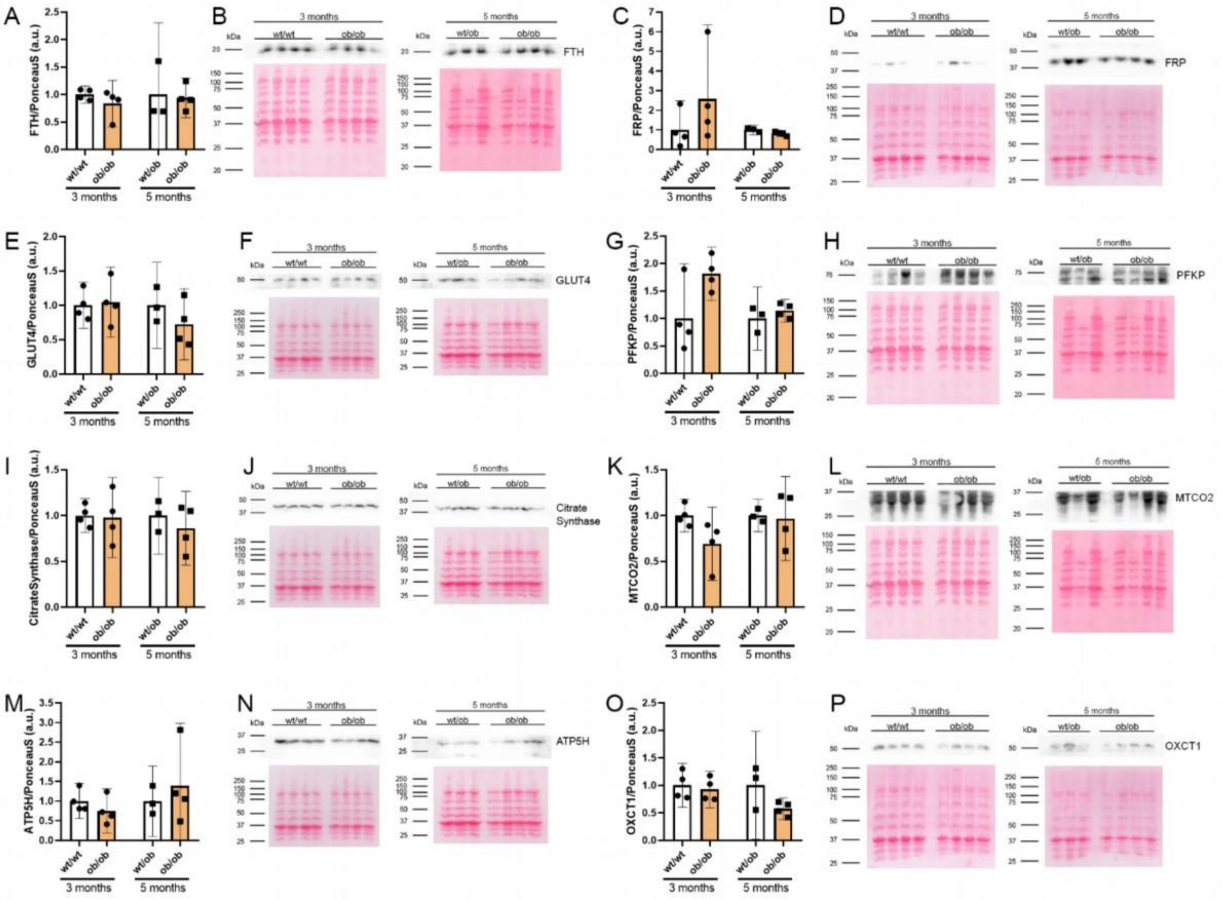


**Supplementary Figure 7. Evaluation of proteins involved in energy metabolism in gastrocnemius muscle.** The relative quantification of ferritin (FTH, **A and B**), ferroportin (FRP **C and D**), GLUT4 (**E and F**), platelet isoform phosphofructokinase (PFKP, **G and H**), citrate synthase (**I and J**), mitochondrially encoded cytochrome C oxidase II (MTCO2, **K and L**), ATPase Subunit D (ATP5H, **M and N**), and 3-Oxoacid CoA-Transferase 1 (OXCT1, **O and P**) were carried out by western blotting. Bars represent the group mean, error bars represent the confidence interval of 95%, and each dot represents a biological sample. Representative images of immunodetection with respective ponceau S staining are presented next to each graph.

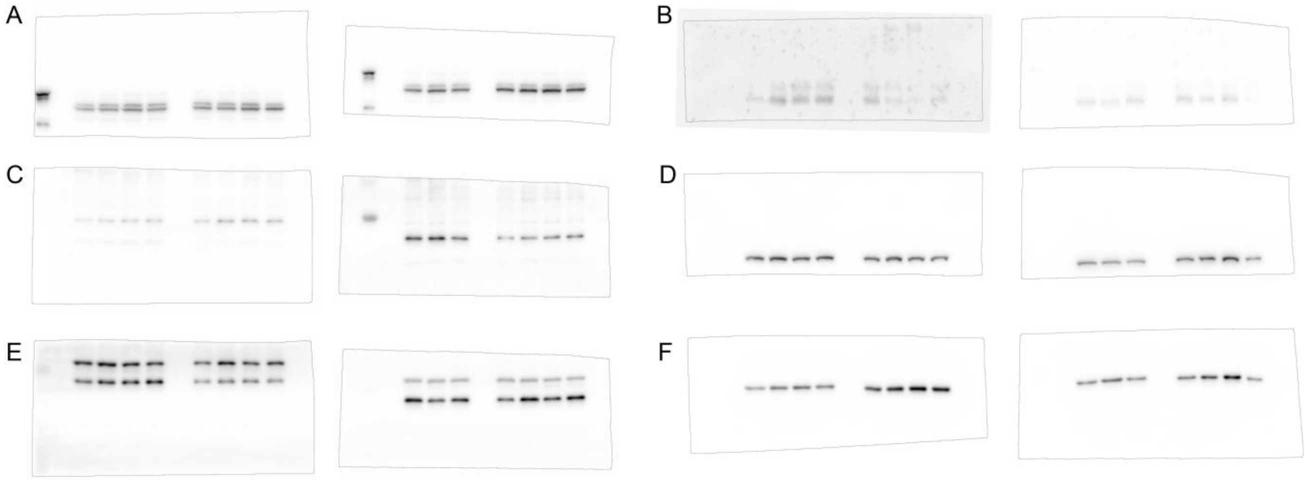


**Supplementary Figure 8. Ferritin and metabolic proteins in the hippocampus.** Uncropped images of Figure 6. **A.** OXCT1. **B.** GLUT1. **C.** Ferritin. **D.** Citrate Synthase. **E.** ATP5H. **F.** MTCO2. The membranes were cut at 37kDa to enable the detection of multiple proteins with distinct molecular weights using the same membrane.

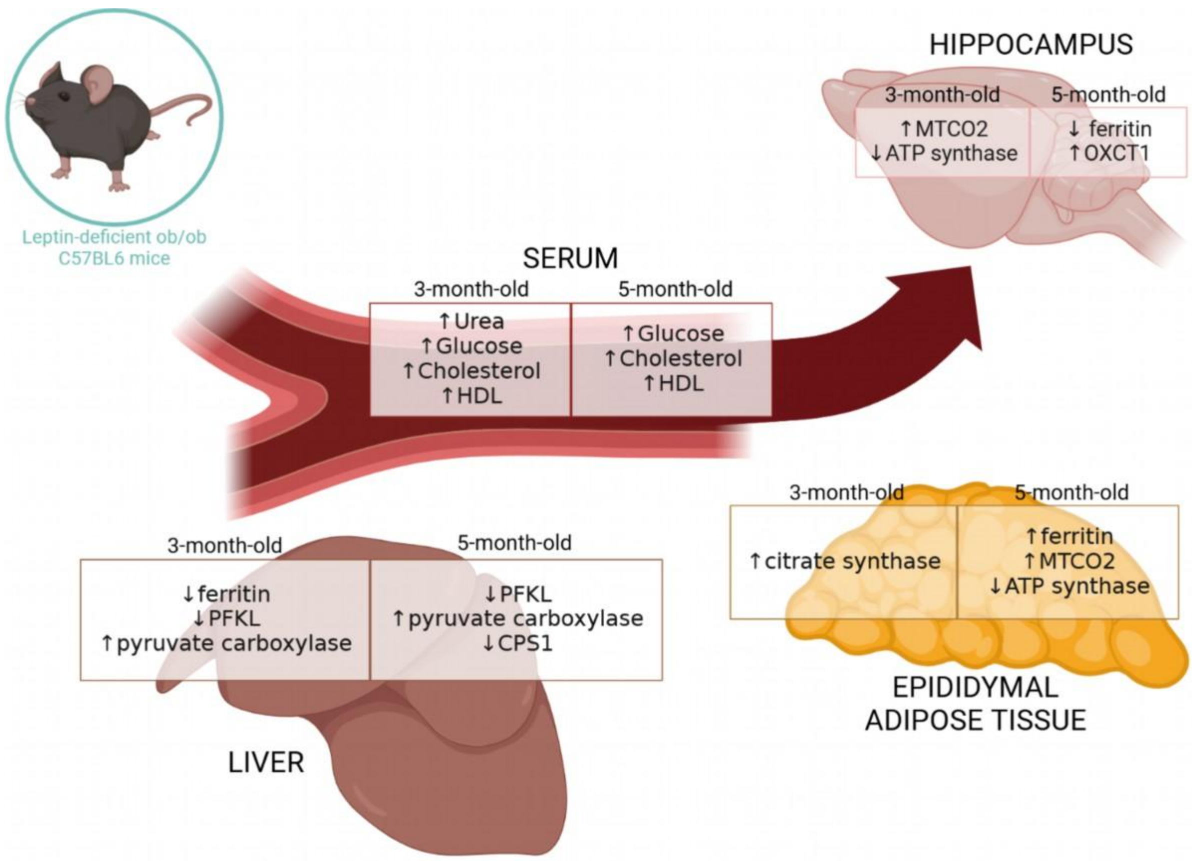


**Supplementary Figure 9 Overview of metabolic changes and alteration of iron homeostasis in multiple organs.** Proteins or metabolites that showed significant changes in ob/ob mice were summarized.
